# Emergent Antibiotic Persistence in Cross-Feeding Bacteria in Structured Habitats

**DOI:** 10.1101/2023.10.09.561574

**Authors:** Xianyi Xiong, Hans G. Othmer, William R. Harcombe

## Abstract

Antibiotic persistence allows a sub-population of bacteria to survive antibiotic-induced killing and contributes to the evolution of antibiotic resistance. Although bacteria typically live in microbial communities with complex ecological interactions, little is known about how microbial ecology affects antibiotic persistence. Here, we demonstrated that the combination of cross-feeding and community spatial structure can emergently cause high antibiotic persistence in bacteria by increasing the cell-cell heterogeneity in the metabolic state both during growth and antibiotic killing. We tracked the ampicillin-induced death on agar surfaces in a model obligate mutualism of *Escherichia coli* and *Salmonella enterica*. We found that *E. coli* formed ∼100-fold more antibiotic persisters in the cross-feeding coculture than in monoculture. This high persistence could not be explained solely by the presence of *S. enterica*, the presence of cross-feeding, or the growth rate differences between the mono- and co-cultures. Time-series fluorescent microscopy revealed increased cell-cell variation in *E. coli* lag time in the mutualistic co-culture. Furthermore, we discovered that an *E. coli* cell can survive antibiotic killing if the nearby *S. enterica* cells on which it relies die first—a mechanism we termed “dynamical loss of access to nutrient.” In conclusion, we showed that the high antibiotic persistence phenotype can be an emergent phenomenon caused by a combination of cross-feeding and spatial structure. Our work highlights the importance of considering spatially-structured interactions during antibiotic treatment and to understand microbial community resilience more broadly.

## Introduction

Understanding the strategies of bacteria to overcome antibiotic stress is a pressing task in the 21^st^ century. Antibiotic tolerance and persistence are two understudied yet important strategies for bacteria to survive antibiotics [1]. While antibiotic tolerance occurs when the entire bacterial population dies more slowly during the antibiotic treatment [2], antibiotic persistence refers to the situation where just a sub-population of bacteria is killed more slowly [3]. Both tolerance and persistence can contribute to the evolution of antibiotic resistance [4,5], a global public health problem that was associated with ∼4.95 million deaths world-wide in 2019 alone [6]. There is an increasing understanding of the mechanisms of antibiotic tolerance [2,4,7,8], but we remain much more ignorant about the mechanisms behind antibiotic persistence [5,9].

Antibiotic persistence is challenging to study because persister cells (the tolerant *sub-population* of cells) are genetically identical to non-persisters and typically exist at a frequency less than 1%. Antibiotic persistence results from phenotypic heterogeneity among cells in a population [10–12], and can arise due to stochasticity in protein activity or gene expression [13,14]. These phenotypic differences cause a fraction of bacterial cells to enter a physiological state that allows them to persist through antibiotic killing and recover after the antibiotic perturbation [5,9,13,15]. However, most research on tolerance and persistence has been done in monocultures in shaken liquid, which ignores the fact that bacteria in nature often live in structured communities with complex ecological interactions [16,17]. How microbial ecology contributes to antibiotic persistence remains largely unknown.

Microbial ecology may be important to antibiotic persistence because ecological interactions among bacterial species can affect responses to antibiotic treatment [18–28]. One common ecological interaction is cross-feeding [25,29–32], which involves bacteria obtaining nutrients from the excretion of other species [33]. Recently, our group showed that cross-feeding can reduce the antibiotic resistance of species down to that of the most susceptible species on which they rely—a mechanism called the “weakest link” hypothesis [18]. Conversely, positive interactions can change the pH of growth media and increase a focal species’ tolerance against an antibiotic [34]. Positive interactions also can increase the minimum inhibitory concentration (MIC) of antibiotics for cells by promoting more efflux activity and reducing intracellular drug concentrations [25]. Finally, positive interactions can lead to increased protection of a sensitive species by increasing proximity to an antibiotic-degrading strain [35]. However, there has been little study of the impact of interspecies interactions on antibiotic persistence.

Spatial structure also influences microbial ecology in ways that may contribute to antibiotic persistence in interacting species. The spatial structure (i.e. the physical arrangement of cells [36]) of a collection of microbes can increase the phenotypic heterogeneity among individual bacteria in a clonal population by generating differences in access to nutrient [37] and can affect the responses to antibiotics [35]. Biofilms generate more persisters as cells at the center of an aggregation are deprived of nutrients and therefore dormant [38,39]. Similarly, recent research [40] showed that an isogenic lawn of *Escherichia coli* in a spatially-structured microfluidic chamber has a higher survival fraction than in a shaken liquid culture after antibiotic killing. Cells that are further from the nutrient source switched from consuming glucose to cross-feeding acetate [40]. Intuitively, spatial structure in cross-feeding mutualisms may lead cells to have differential proximity to a cross-feeding partner and different access to the cross-fed nutrients [41], thereby increasing phenotypic heterogeneity within a population.

Here, we investigated how the combination of spatial structure and cross-feeding can emergently contribute to the antibiotic persistence in bacteria. We focused on an obligate cross-feeding mutualism between *E. coli* and *Salmonella enterica* that exchange methionine and carbon [42–44]. We found that in a spatially-structured environment with ampicillin, the persister frequency of both species was 32∼100-fold higher in the cross-feeding coculture than in their respective monocultures. Interestingly, antibiotic tolerance was not affected. We focused on *E. coli* to understand the mechanism behind this heightened persistence, and developed time-series fluorescent microscopy experiments to study growth and death in spatially-structured environments on a single-cell level. We discovered that the *E. coli* that relies on *S. enterica* can survive if the *S. enterica* cells nearby die first—an ecological mechanism underlying persistence in the spatially-structured mutualism that we termed “dynamical loss of access to nutrient.”

## Materials & Methods

### Bacterial strains and culture media

The *E. coli* strain used in this study is a K-12 BW25113 derivative from the Keio collection carrying a *ΔmetB* mutation (JW3910) [45], a reinserted *lac* operon [42], and a genome-integrated, constitutively expressed cyan fluorescent protein (CFP) gene [43]. The *S. enterica* LT2 strain constitutively expresses a genome-integrated yellow fluorescent protein (YFP) gene [43], and over-excretes methionine due to a base-pair change in *metA* [46] and an IS element inserted in front of *metJ* [47].

The strains form an obligate cross-feeding mutualism in lactose minimal media [42–44] (Fig. 1A and S1A). In lactose minimal media, *E. coli* cross-feeds methionine from *S. enterica* and secretes a waste carbon product that *S. enterica* requires to grow (Fig. 1A). *E. coli* can also grow in monoculture in the lactose minimal media with methionine supplementation, thus maintaining the same physiology as the cross-feeding counterpart [18] (Fig. 1A and Fig. S1B). The *S. enterica* strain can also be studied in monoculture when supplemented with a carbon source like acetate (Fig. S1B) [42–44].

**Fig. 1.**
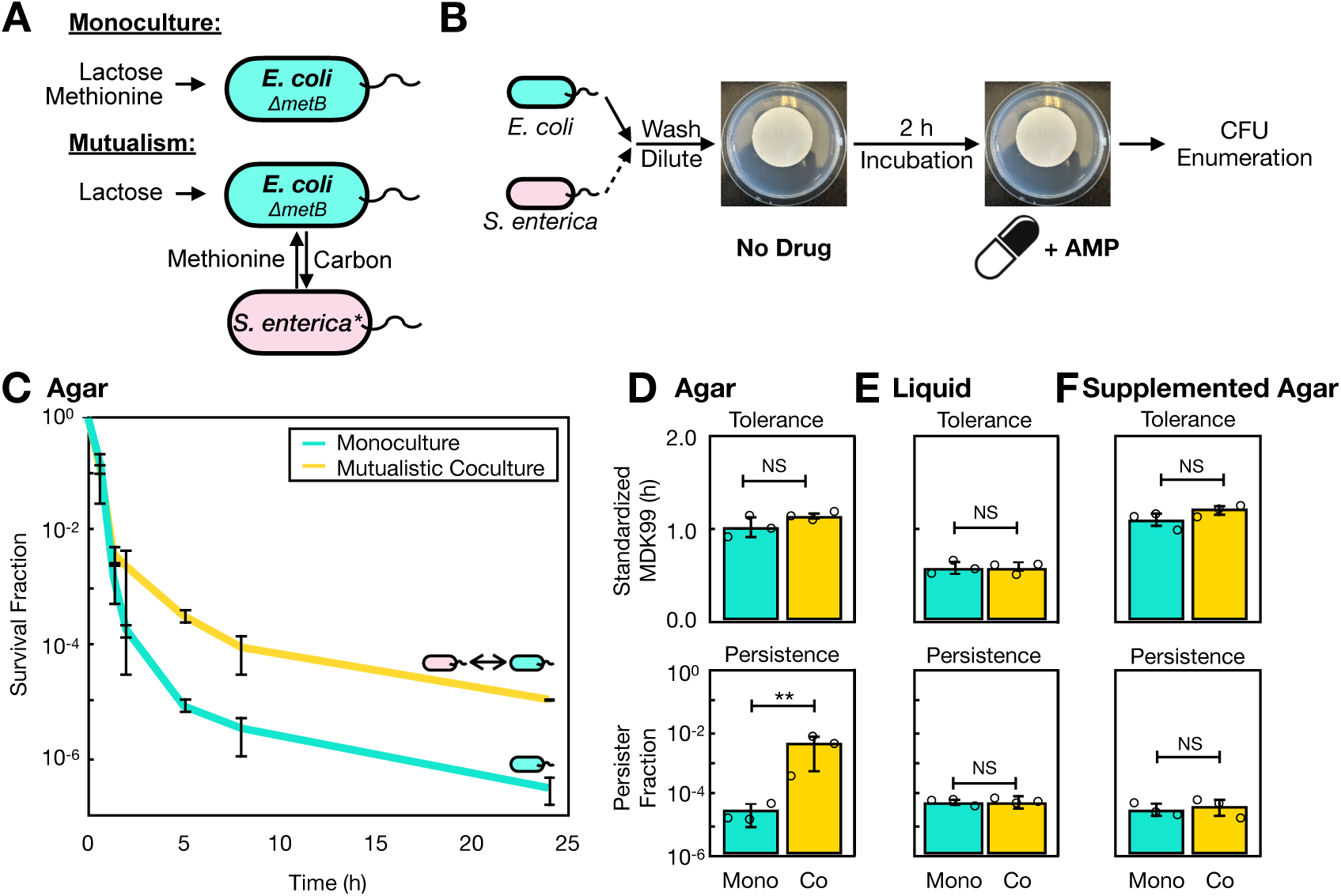
High antibiotic persistence is an emergent phenotype caused by both spatial structure and cross-feeding. **A** Model system setup. An *E. coli* methionine auxotroph can obtain necessary methionine from media (Monoculture) or from a methionine-overproducing *S. enterica* (Mutualism). The coculture represents an obligate mutualism as *S. enterica* relies on *E. coli* for carbon in lactose minimal media. **B** Monocultures or cocultures were immobilized on nitrocellulose filter membranes and incubated on Hypho minimal agar for 2 hours without antibiotics before transferred to agar containing ampicillin (AMP). Membranes were destructively sampled through time to determine survival fractions. **C** *E. coli* survival curves in monoculture and the mutualistic coculture under ampicillin treatment over 24 hours. **D** The antibiotic tolerance and persistence of *E. coli* in monoculture (Mono) and the mutualistic coculture (Co) calculated from **C**. **E** *E. coli* tolerance and persistence in shaken liquid cultures in monoculture (Mono) and the mutualistic coculture (Co). **F** *E. coli* tolerance and persistence in monoculture (Mono) and the methionine-supplemented coculture (Co) on agar. All values shown are averages of measurements from 3 biologically independent trials (*n*=3). Error bars denote standard deviation of these measurements. NS: no significance, *p*>0.05. **: *p*<0.01.

Hypho minimal media was used to study responses to antibiotic killing in *E. coli* and *S. enterica* (see full media recipe in Supplementary Methods 1) [48]. The mutualistic medium contains 2.78 mM lactose, whereas the *E. coli* monoculture or methionine-supplemented coculture medium contains 2.78 mM lactose and 0.08 mM methionine. The *S. enterica* monoculture medium contains 16.9 mM acetate. Agar plates contained 1% (g/mL) agar. When noted in the text, the ampicillin sodium salt (Fisher Scientific, MA) was prepared in the agar or added into the liquid media to reach a final concentration of 100 *μ*g/mL [3], which is >128-fold higher than the MIC of *E. coli* [19].

Fresh (<1 week) *E. coli* and *S. enterica* colonies were inoculated into liquid Hypho minimal media with lactose and methionine, and with acetate, respectively, and incubated with shaking at 37 °C till log phase. *E. coli* and *S. enterica* cells were then washed and diluted in saline [5] to OD_600_=0.005 or OD_600_=0.0025, respectively. For coculture assays, cells were mixed at these OD_600_ to achieve a 1:1 species ratio.

### Studying bacteria on spatially-structured membranes

An amount of 2 mL of the above diluted culture (∼5×10^6^ cells/species/membrane) was then spread evenly and immobilized on a 0.2 *μ*m nitrocellulose filter membrane (4.7 cm diameter; Thermo Fisher Scientific, MA) on a flat-bottom, ethanol sterilized funnel by removing liquid with an air vacuum. In antibiotic killing assays, the membrane was incubated on Hypho agar without antibiotics to allow growth to onset for 2 hours, and then moved to a different agar with 100 *μ*g/mL ampicillin (Fig. 1B and S1C). In growth curve assays, the membranes were incubated on Hypho agar without antibiotics.

To determine population sizes at each time point, we sacrificed a membrane per biological replicate. Membranes were vortexed for 30 s in 5 mL saline to wash off the cells. For ampicillin killing experiments, a stock solution of 125 unit/mL of *β*-lactamase (Neta Scientific, NJ) was added [3]. Dilutions were plated on differential LB agar with 20 *μ*g/mL 5-bromo-4-chloro-3-indolyl-β-D-galactopyranoside (X-gal). After 1 day incubation at 37 °C and 5 days at room temperature (∼ 24 °C), we calculated the population size of *E. coli* and *S. enterica* by enumerating the CFUs (colony forming units) of blue and white colonies, respectively. Note that the long incubation was to ensure that all colonies—including the small colony variants—appear and form visible colonies with proper color [50].

### Growth physiology measurements

To measure growth rate and lag time we determined CFU at different time points and fit an exponential line to the log phase of the growth curve. Growth rate was measured as the slope of the curve. Lag time was estimated as the *x*-axis value of the intersection between the exponential fit line and the initial biomass of *E. coli*.

### Antibiotic tolerance and persistence measurement

Survival curves were plotted using the survival fraction data calculated from CFU counts at each ampicillin-treated time point. We measured the persister fraction by fitting an exponential line to the slower phase of the kill curve and then calculating the *y*-axis intersection of the fitted line (Fig. S2A) [1]. Antibiotic tolerance was first estimated with the MDK99 (<u>m</u>inimum <u>d</u>uration for <u>k</u>illing <u>99</u>% of the population) metric as the time it takes to reduce the population to 1% of its original density[10,45]. We also mathematically derived necessary standardization of the MDK99 measurement if the persister fraction is estimated above 1%, which, by definition, will affect the MDK99 value (Fig. S2B-C; Supplementary Methods 2).

### Time-series fluorescent microscopy

For microscopy, cells were placed on agarose pads. Log-phase *E. coli* and *S. enterica* cultures were washed and diluted to OD_600_=0.05 or OD_600_=0.025, respectively. A volume of 1.5 *μ*L of the diluted culture was added onto a dry Hypho minimal agarose pad (W: 0.4 cm × L: 0.4 cm × H: 0.1 cm) with 1% (m/v) agarose (Sigma Aldrich, MO), and dried at room temperature. The agarose pad was then flipped over onto a microscope slide (ibidi USA Inc., WI) so bacteria were between the pad and the slide. The pads were then incubated for 2 hours at 37 °C in the microscope chamber on the Nikon A1si Confocal and Widefield inverted microscope in the University Imaging Centers at the University of Minnesota (St. Paul, MN). For antibiotic experiments, a drop of 1.5 *μ*L ampicillin solution was added on top of the agarose pad on the microscope before imaging such that the pad obtained a drug concentration of 100 *μ*g/mL.

We programmed the Nikon Elements v5.41 software to collect fluorescent signals with 200 total magnification at the center of each agarose pad. The images were taken every 20 min for a total of 7 hours at two fluorescent channels: 488 nm (CFP) for *E. coli* and 514 nm (YFP) for *S. enterica*. The collected time-series images were then aligned and noise was removed using the software. In particular, a rolling-ball algorithm (radius: 1.27 *μm*) was implemented around each bright object to subtract background. An automatic deconvolution method was used to enhance contrast and remove blur (modality: Point scan confocal; pinhole size: 93.23 *μ*m; magnification: 20.0 ×; numerical aperture: 0.75; immersion reference index: 1.0 [air]; calibration: 0.156 *μ*m/pixel). Local contrast was set at 25% for a radius of 2.34 *μ*m per object. Single cells were identified by detecting regional maxima at the center from a 5×5 matrix. The fluorescent signal threshold was set to 100 unit of intensity to produce binary images of individual cells, which were objects that passed the threshold. Cells on the image borders were also removed from analysis. Images were then further analyzed using ImageJ (FIJI, v1.53k) [51].

For the growth analysis images were in a dimension of 160 *μ*m × 160 *μ*m. There were usually ∼100 cells per image. ImageJ was used to identify cell clusters as cells localized next to each other that the image processing steps above could not separate. We then took the images from the first 4 hours for analysis because clusters started to fuse together after that time due to growth. The cell cluster biomass was defined as the fluorescent signal area per cluster. We identified the minimal time frame at which each cluster gains a biomass of 10%, which we considered lag time in a similar process following previous work [2,7,36,52].

For the death analysis images were in a dimension of 476 *μ*m 476 *μ*m, which is larger than in the growth analysis as persisters usually exist in low frequency so more cells have to be observed. There were usually 500∼700 cells per image frame. In the ampicillin treatment, we rarely observed clusters of more than 2 cells primarily due to less division under the antibiotic stress. We used ImageJ to measure single object area (biomass) at each location over time. Objects were considered cells when >25 pixels of the respective fluorescent signals were detected. We then considered the single-cell death time as the earliest time frame for a cell to disappear from the microscopic view.

### Statistics and data availability

One-way analysis of variance (ANOVA) tests and Spearman’s correlation tests were implemented in R v3.3.3 [53]. For multiple comparisons tested with one-way ANOVA, we reported *p*-values adjusted with Tukey’s HSD (honestly significant difference) [54]. All source data were attached as part of the Supplementary Information of this work.

## Results

### Spatially-structured cross-feeding causes heightened antibiotic persistence

To investigate whether cross-feeding affects bacterial response to antibiotic killing on spatially-structured surfaces, we incubated *E. coli* in monoculture and in the cross-feeding coculture with *S. enterica* on agar plates and measured its survival curves when treated with high-dosage ampicillin (Fig. 1A-B and S1C; Materials & Methods). We found that *E. coli* had different, biphasic survival curves in monoculture and in mutualism (Fig. 1C). Using these survival curves, we estimated the *E. coli* antibiotic tolerance and persistence. Although *E. coli* has a similar antibiotic tolerance in mutualism and in monoculture (*p*=0.2), the mutualistic *E. coli* on average has a ∼100-fold higher antibiotic persister frequency in coculture (*p*=0.009; Fig. 1D). We also observed similar tolerance but higher persistence in *S. enterica* in the mutualistic coculture relative to monoculture (Fig. S3). Together, these results suggested the generality of our observation that cross-feeding in a spatially-structured habitat can increase antibiotic persistence.

To further understand our above observation, we focused on *E. coli* and tested whether cross-feeding is sufficient to drive the high antibiotic persistence. We repeated the experiment in shaken liquid without spatial structure [40] and maintained similar average cell-cell distances as on surfaces (Fig. S4A-B; Supplementary Discussion 1). We found that in liquid cultures with ampicillin, *E. coli* has similar antibiotic tolerance (*p*=0.9) and persistence (*p*=0.7) in monoculture and the mutualistic coculture (Fig. 1E). Lowering the culture density in the liquid mutualistic coculture up to ∼100 fold did not cause the high persistence observed in the structured cross-feeding (Fig. S4).

We next examined whether cross-feeding is essential for the high *E. coli* persistence. We broke the metabolic dependency of *E. coli* on *S. enterica* by supplementing methionine in the agar, and observed that *E. coli* does not show the high persistence phenotype without cross-feeding with *S. enterica* for methionine (*p*=0.7; Fig. 1F). These data suggested that the presence of *S. enterica* is not enough to cause more *E. coli* persisters in structured habitats, and that cross-feeding between the two species was also a necessary factor.

### Population growth dynamics do not explain the high persistence when cross-feeding

Slower growth or longer lag time are associated with persistence [2], so we decided to study the *E. coli* growth physiology in the spatially-structured mutualism. We counted CFUs on membranes to obtain the *E. coli* growth curves (Fig. S5) and calculated growth rates and lag times (Materials & Methods). We found that on surfaces, *E. coli* on average grows more slowly in the cross-feeding coculture than in monoculture (Tukey’s HSD *p*=0.006) but the mean lag times are similar (Tukey’s HSD *p*=0.9; Fig. 2A). To test whether the slower average growth rate in mutualism is a sufficient driver for more persisters, we lowered the monoculture *E. coli* growth rate by reducing temperature to 30 °C, and showed that *E. coli* on average grows at a similar rate in the 37 °C mutualism as in the 30 °C monoculture (Tukey’s HSD *p*=0.9; Fig. 2A). However, lowering growth rate in the *E. coli* monoculture did not increase tolerance (Tukey’s HSD *p*=0.2) or persistence (Tukey’s HSD *p*=0.2; Fig. 2B). These data suggested that the average growth rate differences between monoculture and mutualism are not sufficient to affect the *E. coli* tolerance or persistence on surfaces.

**Fig. 2.**
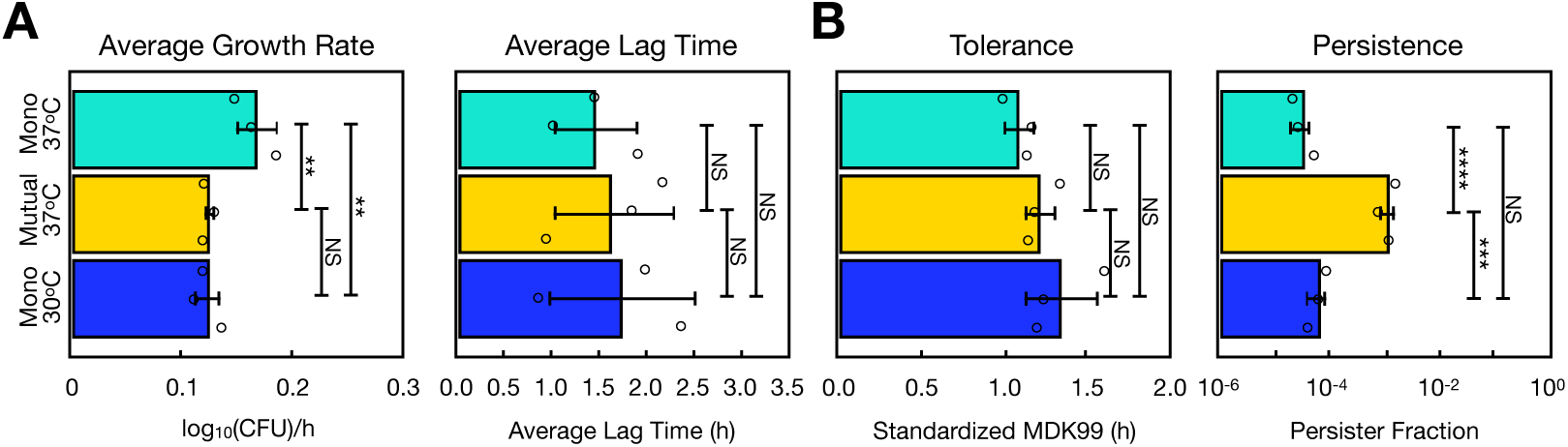
Average *E. coli* growth rate differences in monoculture and mutualism do not explain the persistence difference. **A** The average *E. coli* growth rate and lag time was assessed for populations grown on nitrocellulose membranes at 37 °C and 30 °C in monoculture (Mono) and the mutualistic coculture (Mutual). **B** The antibiotic tolerance and persistence were also measured. All values shown are average measurements from 3 biologically independent trials (*n*=3). Error bars denote standard deviation of these measurements. NS: no significance, *p*>0.05. ***: 1E-4<*p* 1E-3. ****: 1E-5<*p* 1E-4.

We also tested whether starvation [5] of methionine contributes to more *E. coli* persisters. While starvation increased the number of persisters as previously shown [5], it also dramatically increased the antibiotic tolerance (Fig. S6; Supplementary Methods 2). Taken together, the data ruled out most average population-level differences between the monoculture and mutualistic *E. coli* as major contributors to the high persistence phenotype. These results also suggested that the high persistence in *E. coli* is an emergent property caused by the combination of cross-feeding and spatial structure.

### Cross-feeding in structured habitats increases lag time heterogeneity in *E. coli*

We hypothesized that cross-feeding and spatial structure together increase the cell-to-cell heterogeneity in the *E. coli* lag time during growth, but not the average lag. We used time-series fluorescent microscopy to directly observe *E. coli* on a single-cell level while it grows on spatially-structured agarose pads [55], and measured lag time for single cell clusters (Fig. 3A and S7; Materials & Methods). We confirmed that there is a higher variation in the *E. coli* lag time in the cross-feeding mutualism than in monoculture (*p*=0.03; Fig. 3B). This result is also supported by a mathematical model on individual colony growth (Fig. S8 and Supplementary Methods 3). Consistent with our population-level lag time measurement (Fig. 2A), the average lag time measured from the microscopic *E. coli* clusters is similar between monoculture and mutualism (*p*=0.2; Fig. 3B). Together, these data encouraged us to ask why there is higher lag time variation in the mutualistic coculture.

**Fig. 3.**
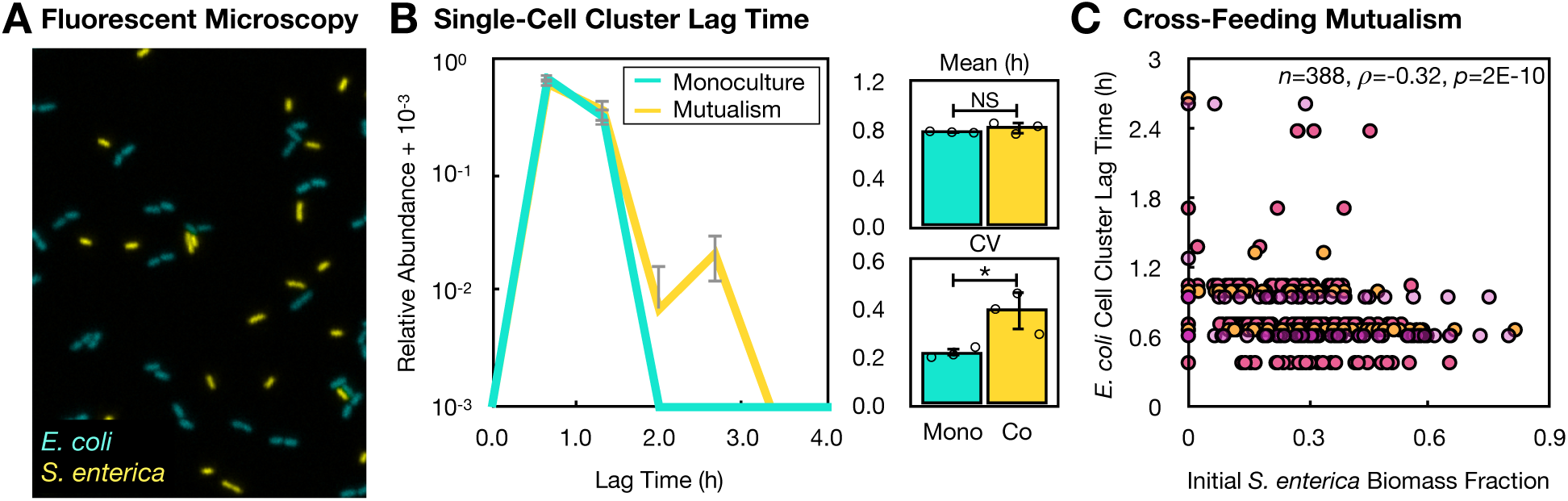
cross-feeding and spatial structure increase cell-cell heterogeneity in the *E. coli* lag time during growth. **A** A time-series fluorescent microscopy experiment was done to track the growth for each single *E. coli* cell cluster (Supplementary Methods). **B** The distribution of the *E. coli* lag time measured in monoculture (Mono) or in the mutualistic coculture (Co) on Hypho minimal agarose pads. The average and coefficient of variation (CV) of lag time values are shown from three independent experiments (*n*=3). Error bars denote standard deviation. NS: no significance, *p*>0.05. *: *p*<0.05. **C** The initial biomass fraction of *S. enterica* within a small (radius = 8.6 m) neighborhood of each *E. coli* cell cluster can predict the lag time of *E. coli* clusters. Each color represents data from one of the three biological replicates (*n*=3).

We proposed that the spatial structure of the cross-feeding mutualism determines the *E. coli* lag times. Previous research on the *E. coli-S. enterica* cross-feeding system suggested that on surfaces, each *E. coli* cell likely interacts with *S. enterica* in local neighborhoods within a short range [42–44]. To experimentally confirm this, we tested whether each *E. coli*’s lag time can be predicted by its proximity to the cross-feeding *S. enterica* partners. We validated that in a neighborhood with a radius of ∼10 *μ*m around each *E. coli* cluster, higher fractions of the *S. enterica* biomass in the beginning of the experiment correlate with shorter *E. coli* lag times (Spearman’s *ρ*=-0.316, *p*=2E-10; Fig. 3C). Furthermore, the above correlation becomes weaker as we considered larger neighborhood sizes (Fig. S9), supporting previous notions that microbial mutualisms typically interact within short ranges [42–44,56]. Together, we concluded that the initial spatial distribution of cells in the cross-feeding mutualism can explain the increased lag time heterogeneity, which may be key to having more persisters.

### Fluorescent microscopy revealed an unexpected mechanism behind high *E. coli* persistence in the cross-feeding mutualism

Finally, we established a time-series fluorescent microscopy protocol on spatially-structured agarose pads to measure the death time of single *E. coli* cells in the presence of ampicillin (Fig. S10; Materials & Methods). We found that cross-feeding increased the variance (as coefficient of variation or CV, *p*=0.02) but not the mean death time (*p*=0.09) relative to the monoculture counterpart (Fig. 4A). This finding is consistent with higher persistence but similar tolerance in the spatially-structured mutualism. We found that the individual *E. coli* death time did not correlate with the initial fraction of *S. enterica* biomass around each *E. coli* cell or the initial distance to the nearest *S. enterica* neighbor (Average Spearman’s *ρ*=0.00661±0.0509, *n*=3; Fig. 4B). Thus, although the initial spatial structure of the mutualism is sufficient to predict lag time (Fig. 3C), it is insufficient to explain the higher persister fraction.

**Fig. 4.**
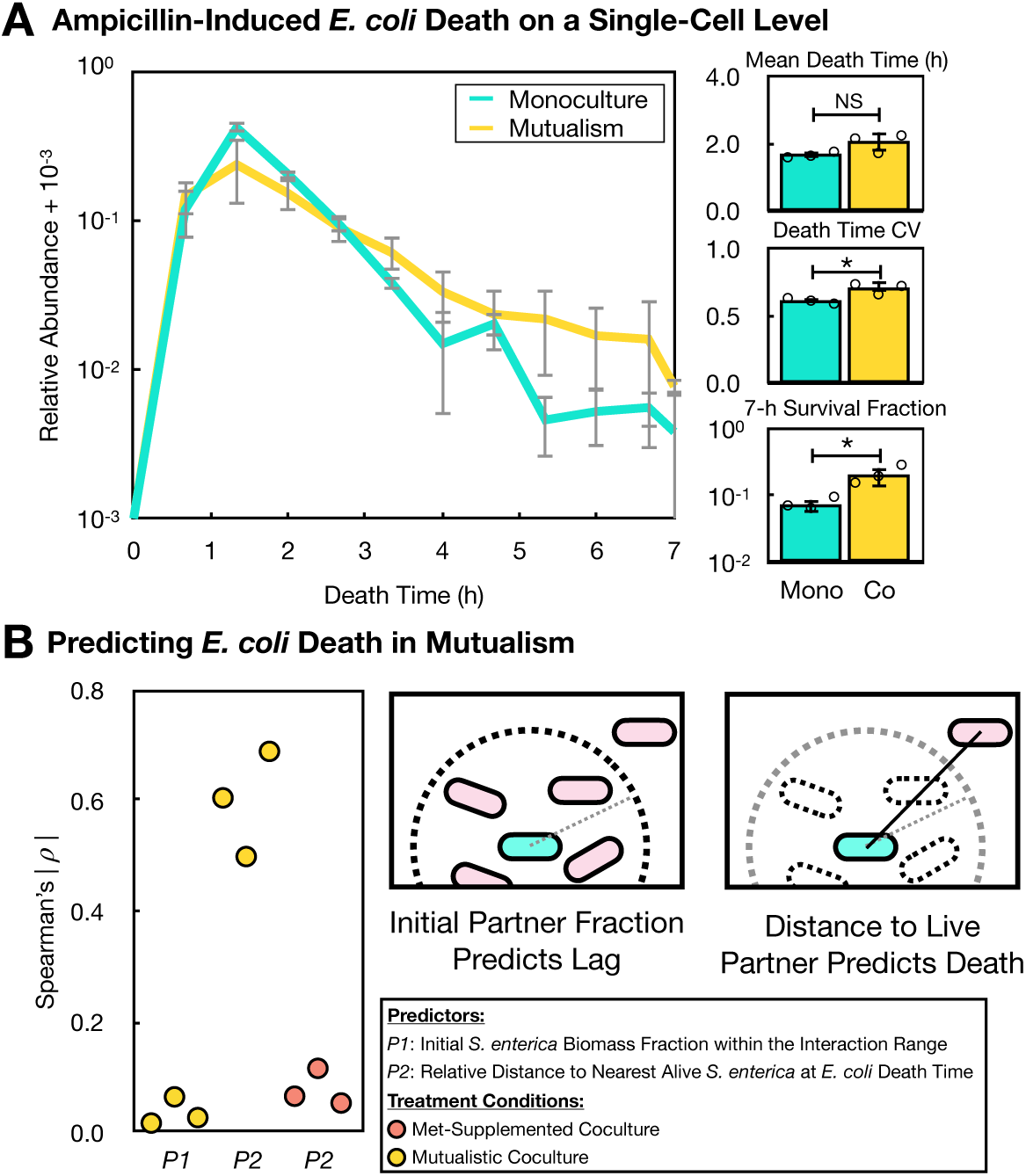
Dynamic loss of access to nutrient (DLAN) underlies the higher *E. coli* persistence in the structured mutualism. **A** The death time distributions of *E. coli* during ampicillin treatment within 7 hours of imaging. The mean, CV of single-cell death time for *E. coli* that die within 7 hours, and the survival fractions after 7-hour killing were calculated for the monoculture (Mono) and the mutualistic coculture (Co). Error bars denote standard deviation of the average measurements from 3 biologically independent trials (*n*=3). NS: no significance, *p*>0.05. *: *p*<0.05. **B** The absolute Spearman’s in each biological replicate was plotted for Predictors (*P1*, *P2*) of the single-cell *E. coli* death time in the mutualistic or the methionine- (Met-) supplemented coculture. Schematic: cyan cell: live *E. coli*; pink cell: live *S*. *enterica*; dashed cell: dead *S. enterica*; dashed circle: interaction range; dashed line: radius of interaction range; black line: distance to live *S. enterica*.

Next, we tested whether the dynamic change of spatial structure in mutualism over time determines the *E. coli* single-cell death time. Because *S. enterica* are killed by ampicillin as well, perhaps it is crucial to consider proximity over time rather than just the initial proximity of *E. coli* to its partner. At the time of death for each *E. coli* cell, we determined the distance to the nearest alive *S. enterica*. We then standardized this distance against the average distance between partners within the microscopic frame at each time point (Supplementary Discussion 1), in order to prevent artificial correlations when fewer cells are present in the microscopic frame at later time points. We found that taking *S. enterica* death into account in this way leads to a much stronger correlation between proximity and death time (Average Spearman’s *ρ*=0.598±0.0953, *n*=3; Fig. 4B). Thus, this result confirmed a strong spatial pattern contributing to the increased *E. coli* persistence in the mutualism.

To further validate our metric, we tested the predictive ability when *E. coli* is challenged with ampicillin in coculture but not cross-feeding with *S. enterica*. We found that in the methionine-supplemented facilitative coculture, the *E. coli* death time does not correlate with the standardized distance to the alive *S. enterica* neighbor (Fig. 4B; Average Spearman’s *ρ*=0.0790±0.0330, *n*=3). Together, these findings suggested that the temporal dynamics of the mutualistic community spatial structure determines the *E. coli* death time throughout the ampicillin treatment. *E. coli* cells lose access to methionine as their *S. enterica* neighbors die and this loss increases *E. coli*’s chance of survival in the ampicillin treatment (Fig. 4B). We call this mechanism “dynamical loss of access to nutrient” (DLAN) to reflect the mutualistic nature of the involved interspecies interactions.

## Discussion

In this work, we demonstrated that cross-feeding in a spatially-structured environment can increase antibiotic persistence by 32∼100-fold in both the mutualistic *E. coli* and *S. enterica* as compared to the monocultures. We found that the high persistence phenotype in *E. coli* is an emergent property that only appears when cross-feeding takes place in a structured habitat, and showed that many population-level physiology differences between monoculture and mutualism (e.g. the average growth rate) are not sufficient to explain the increase in persistence. Using time-series fluorescent microscopy to directly observe individual *E. coli* cells, we found a higher variation in the lag time in the mutualistic coculture that can be attributed to the initial proximity to mutualistic partners. However, this initial spatial structure cannot predict the single *E. coli* cell death time, suggesting a previously uncharacterized ecological mechanism for persistence. By directly observing the ampicillin-induced death in *E. coli* and *S. enterica* cells, we discovered that the *E. coli* that relies on *S. enterica* can survive the antibiotic killing if the *S. enterica* cells nearby die first—a mechanism we named “dynamical loss of access to nutrient.”

A key discovery in our work is that cross-feeding in structured habitats can result in more persisters. Antibiotic persistence has been shown to be the result of cell-to-cell variation in physiology in a population [3,5,12–15]. This variation can arise from stochastic differences in transcription [14] or translation [13] levels among cells. Our observation suggests that cross-feeding on a surface introduces additional cell-to-cell variation as a result of stochastic differences in how close a cell lands to its cross-feeding partners. In liquid cultures nutrients diffuse rapidly, reducing environmental heterogeneity and the stochastic effects of location. High antibiotic persistence is therefore an emergent property resulting from the interplay between two important ecological factors, interspecies interactions and spatial structure (Fig. 1).

Variation in lag time was well predicted by the frequency of partners in a small neighborhood (Fig. 3). Growth on surfaces typically makes interactions local [37,42,43,44], and our results suggest that cross-feeding is strongest among cells within ∼10 *μ*m in the *E. coli*-*S. enterica* system. This result is in agreement with previous work [56] suggesting that growth of cross-feeding *E. coli* is predicted by the fraction of partners within 3∼12 *μ*m. These results highlight that the location of bacteria is critical for understanding which cells interact in microbial systems.

The lag time variation caused by initial partner proximity was insufficient to predict the *E. coli* death time (Fig. 4B). Death of cells quickly alters the proximity of cross-feeding partners, so it is critical to account for this dynamic loss of access to nutrient (or DLAN; Fig. 4B). We would have missed this critical mechanism if we assumed that the sub-population behavior in bacterial cells during growth can always predict their death patterns as in previous research [2,5,7,9,14,15,40,57]. A few studies [3,5,14,40] did directly observe the death time of individual bacterial cells in antibiotics. Studies [3,40] that have measured death times directly do often find correlation between growth and death suggesting that the lack of correlation we observed may be the result of interspecies interactions. Strikingly, our microscopy experiment (Fig. 4) suggests that the cross-feeding *S. enterica* cells must continually produce methionine to support the growing *E. coli* cells in the neighborhood. This result emphasizes that location is not a static feature of microbial communities but rather a dynamic attribute that changes with birth and death of cells.

The current work indicates that the antibiotic persistence in human-associated microbes may be higher than monitored in clinical laboratories [58]. Lab studies often exclude the spatially-structured interspecies interactions that are prevalent in the human microbiome [59]. There is growing appreciation that cross-feeding may be more common in the human body than previously thought (e.g. *Pseudomonas aeruginosa* cross-feeds with the mucin-degrading anaerobes in the Cystic Fibrosis lungs [60]). While previous work assumed that mutualism or spatial structure is not common in the gut [61,62], recent evidence showed that the mammalian gut microbiomes are highly structured [63–65] especially on the mucus layer [66]. Furthermore, consuming human therapeutic drugs can induce cross-feeding in the gut microbiota [67]. Our results thus suggest that antibiotic persistence in the human-associated microbiota is underestimated.

Our current research also has implications for the evolution of antibiotic resistance. Previous *in vitro* [7] and clinical [4] studies found that antibiotic tolerant or persistent bacteria evolve antibiotic resistance faster. Our current results suggest that spatially-structured mutualisms will evolve resistance faster than the monoculture counterparts during antibiotic treatment. This deduction challenges our group’s previous discovery [19] that cross-feeding in shaken liquid cultures slows the evolution of antibiotic resistance, potentially because that study [19] lacked spatial structure and did not cyclically expose bacteria to high-concentration antibiotics as in the clinical reality [2].

In the current study, we show that microbial ecology can affect antibiotic persistence. Our work highlights that microbial interactions and spatial structure can generate emergent increase in individual heterogeneity. As we work to develop precision management of human microbiomes, it will be critical to continue to advance our understanding of the scale over which interactions occur, and how these interactions shape the behavior of individual cells, populations, and communities.

## Acknowledgement

We acknowledge that the microscope experiment was conducted on the Nikon A1 microscope at the University of Minnesota Imaging Center with training and image analysis help from M. Sanders and M. E. Brown. We thank J. M. Chacón, A. Bisesi, J. N. V. Martinson, J. A. Gralnick, M. F. Freeman, S. Ishii (U of MN), J. A. Lee, N. Moreno, J. L. Gonzalez (NASA), C. Song (Princeton), K. R. S. Hale (U of MI), M. Dal Bello, H. Lee (MIT) for the helpful feedback.

## Author Contributions

X.X. and W.R.H. conceived and designed the study. X.X. collected data, performed analysis, and wrote the manuscript. H.G.O. and W.R.H. supervised the project. All authors contributed to the manuscript editing. This work was developed from the master’s dissertation of X.X. at the University of Minnesota [68].

