## Supplementary Materials for "Emergent Antibiotic Persistence in Cross-Feeding Bacteria in Structured Habitats"

**This PDF file includes:**

Supplementary Methods 1-4
Supplementary Discussion 1-2
Supplementary Figures S1-S11
Supplementary Table 1
Supplementary References

For source data see attached Excel file "Xiong_Raw_Data.xlsx"

### Supplementary Methods

#### 1. Culture media

Hypho Minimal Medium: Hypho minimal medium contains 7.26 mM $K_2HPO_4$, 9.38 mM $NaH_2PO_4$, 1.89 mM $(NH_4)_2SO_4$, 0.41 mM $MgSO_4$, 0.6 µM $ZnSO_4$, 9.98 µM $CaCl_2$, 0.5 µM $MnCl_2$, 1 µM $(NH_4)_6Mo_7$, 0.5 µM $CuSO_4$, 1 µM $CoCl_2$, 0.169 µM $Na_2WO_4$, 8.88 µM $FeSO_4$ (Delaney et al., 2013). The mutualistic coculture medium contains 2.78 mM lactose as the sole carbon source (Fig. S1A). In the *E. coli* monoculture, the media and agar plates both contain 2.78 mM lactose and 0.08 mM methionine (Fig. S1B). The *S. enterica* monoculture medium contains 16.9 mM acetate or 5.84 mM galactose as the sole carbon source as specified in the text (Fig. S1B). For the antibiotic killing experiments, the ampicillin sodium salt (Fisher Scientific, MA) was added into the media or prepared within the agar at a concentration of 100 µg/mL (Balaban et al., 2004; >128x of the minimal inhibitory concentration of our *E. coli* strain). When appropriate, Hypho minimal agar was prepared by adding 1% (g/mL) of agar powder (Sigma Aldrich, MO) to distilled water to be autoclaved, and the filter-sterilized carbon source, salts, and ampicillin were aseptically added.

LB Medium: LB agar was made for enumeration of CFU (colony forming units). LB agar was made by adding 10 g agar, 10 g NaCl powder, 5 g yeast extract, and 10 g tryptone powder into 1 L DDI (distilled and deionized) water prior to autoclave sterilization. X-gal (5-bromo-4-chloro-3-indolyl-β-D-galactopyranoside) was always added at 20 $\mu$g/mL into the post-autoclave LB agar medium to distinguish between the lactose-metabolizing *E. coli* (blue colonies) and the non-lactose-metabolizing *S. enterica* (white colonies).

#### 2. Antibiotic tolerance & persistence assays

Antibiotic Persistence Measurement: Following Balaban et al. (2019), we estimated persister fraction by fitting an exponential curve to the less steep (second) phase of each survival curve (e.g. Fig. 1C). This was to emphasize that persisters are a subpopulation of slower-dying cells. The intersection between the less steep exponential curve with the *y*-axis was the considered persister fraction (Fig. S2A).

Antibiotic Tolerance Measurement: As in Fridman et al. (2014) and Brauner et al. (2017), tolerance can be measured as the minimal duration of killing 99% of the population (MDK99) by fitting an exponential line between the two close-by time points that cover the Survival Fraction = $10^{-2}$ data point.

However, by definition, this metric fails to produce reliable tolerance measurements when the persister fraction is higher than 1% (Fig. S2B-C; Supplementary Methods 4). Thus, we measured tolerance for the non-persister population as the "Standardized MDK99" by considering the measured persister fraction (see below). If the persister frequency is measured as $p$ with an experimentally measured MDK99 of $M$, then the population tolerance is defined as:

$$\text{Tolerance} = \begin{cases} M & p \leqslant 10^{-2} \\ M \cdot \dfrac{f\left(10^{-4}\right)}{f(p)} & p > 10^{-2} \end{cases}, \quad [S0]$$

where $f\ (p)$ is the function in Fig. S2C that maps a persister fraction, $p$, to the theoretical MDK99 measurement with no change to the non-persisters' tolerance.

#### 3. PDE-based biophysical model of bacterial colony growth

Model Setup: We developed a mathematical model using partial differential equations (PDEs) to computationally simulate the growth of individual *E. coli* and *S. enterica* colonies in monoculture and the mutualistic coculture on a 1-dimension line.

In the resource-explicit model, bacterial colonies interact with local nutrients in a spatially-structured environment to grow. On a 1-dimension line of distance $Y = 5$, individual cells of *E. coli* (*E*) and/or *S. enterica* (*S*) were seeded randomly by setting the initial conditions at $t = 0$ to be non-zero at random locations, *x*, whereas various nutrients were initially overlaid evenly on the entire line. This random seeding of bacterial cells makes the PDE model able to describe bacterial populations and communities with spatial structure. Each PDE in Equations [1~2] denote the rules regarding how abundance of bacteria or nutrients changes over 40 units of time, $t \in [0,40]$, at any particular *x*.

Equations in [S1] denote how *E. coli* and *S. enterica* cross-feed and grow in a mutualistic community. First, the growth of *E. coli* and *S. enterica* follow Eqs. [S1a] and [S1b], respectively. *E. coli* grows by consuming methionine (*M*) and lactose (*L*), whereas *S. enterica* grows by taking in acetate (*A*). The maximum growth rates are $r_E$ and $r_S$, respectively, and the instantaneous growth rates are limited by the nutrient's local concentration in a Monod fashion with the half-saturation constants, $K_M$, $K_L$, and $K_A$. Growth is ultimately limited by exhaustion of nutrient explained below. *E. coli* growth rates depend on both methionine and lactose, while *S. enterica* growth rates depend only on acetate (see Model Assumptions).

$$\frac{\partial E}{\partial t} = r_E \left(\frac{M}{M + K_M}\right)\left(\frac{L}{L + K_L}\right) E - \kappa_E E. \qquad \text{[S1a]}$$

$$\frac{\partial S}{\partial t} = r_S \left(\frac{A}{A + K_A}\right) S - \kappa_S S. \qquad \text{[S1b]}$$

Eq. [S1c] denotes the temporal dynamics of methionine. As a nutrient source, methionine diffuses in space over time at a rate, $D_M$. It is produced by *S. enterica* at a rate, $p_M$, and is proportional to *S. enterica*'s consumption of acetate in a Monod fashion (Hammarlund et al., 2021). In another word, *S. enterica* only produces methionine when metabolizing on acetate. Methionine gets consumed by *E. coli* following a Monod kinetics by a maximum rate, $c_E$. Finally, methionine degrades slowly over time at a rate, $\kappa_M$.

$$\frac{\partial M}{\partial t} = D_M \nabla^2 M + p_M r_S \left(\frac{A}{A + K_A}\right) S - c_E \left(\frac{M}{M + K_M}\right)\left(\frac{L}{L + K_L}\right) E - \kappa_M M. \qquad \text{[S1c]}$$

Similar to methionine, acetate diffuses at a rate, $D_A$, gets consumed by *S. enterica* following Monod kinetics, decays exponentially by $\kappa_A$ (Eq. [S1d]). Note that acetate is produced by *E. coli* at a rate, $p_A$, as both lactose and methionine are being consumed. Acetate concentration decreases due to *S. enterica* consumption by a maximum rate, $c_S$.

$$\frac{\partial A}{\partial t} = D_A \nabla^2 A + p_A r_E \left(\frac{M}{M + K_M}\right)\left(\frac{L}{L + K_L}\right) E - c_S \left(\frac{A}{A + K_A}\right) S - \kappa_A A. \qquad \text{[S1d]}$$

Finally, the temporal dynamics of lactose concentration—which is only consumed by *E. coli*—follows Eq. [S1e]. Here, the natural decay rate of lactose is, $\kappa_L$.

$$\frac{\partial L}{\partial t} = D_L \nabla^2 L - c_L \left(\frac{M}{M + K_M}\right)\left(\frac{L}{L + K_L}\right) E - \kappa_L L. \qquad \text{[S1e]}$$

The Neumann boundary conditions are:

$$\frac{\partial N}{\partial x} = 0 \text{ at } x = 0,\ 5, \qquad \text{[S1f]}$$

for $N \in \{E, S, M, A, L\}$.

To simulate *E. coli* grown in monoculture, we remove *S. enterica*- and acetate-associated terms from Eqs. [1] to form Eqs. [2] below, as methionine and lactose are now supplemented in the growth medium for monoculture *E. coli* and so no production terms are necessary. All variables are identical to those in Eqs. [1].

$$\frac{\partial M}{\partial t} = D_M \nabla^2 M - c_E \left(\frac{M}{M + K_M}\right)\left(\frac{L}{L + K_L}\right) E - \kappa_M M, \quad \text{[S2a]}$$

$$\frac{\partial E}{\partial t} = r_E \left(\frac{M}{M + K_M}\right)\left(\frac{L}{L + K_L}\right) E - \kappa_E E, \quad \text{[S2b]}$$

$$\frac{\partial L}{\partial t} = D_L \nabla^2 L - c_L \left(\frac{M}{M + K_M}\right)\left(\frac{L}{L + K_L}\right) E - \kappa_L L. \quad \text{[S2c]}$$

Similarly, the boundary conditions are:

$$\frac{\partial N}{\partial x} = 0 \text{ at } x = 0,\ 5, \quad \text{[S1f]}$$

for $N \in \{E, M, L\}$.

Model Assumptions: Our PDE models above are based on the following assumptions.

1. Bacterial growth can be achieved by only consuming among 3 types of nutrients (methionine, lactose, and acetate). In experiments, more nutrients sources have to be present to support growth. For example, ammonia is present in the medium for both monoculture and mutualistic coculture. But, the impact of these other nutrient was factored into the growth rate variables $r_E$ and $r_S$.

2. Nutrient production of both *E. coli* and *S. enterica* only occurs when their growth nutrients are present. In reality, *E. coli* initiates the mutualism by producing acetate without having to metabolize on methionine (unpublished data). This assumption allows us to keep the model simple without having to make special mathematical treatment for *E. coli* and is also a convention in modeling the cross-feeding growth (e.g. Harcombe et al., 2014; Hammarlund et al., 2021).

3. Bacterial growth leads to no diffusion of biomass. In reality, bacterial growth on agar surfaces results in colony formation and colony size expansion (Chacón et al., 2018). This can change the average distance among bacterial colonies.

4. The growth rate of each colony reflects the instantaneous growth rate of individual cells on the nitrocellulose membrane before ampicillin treatment in the experiment.

5. Bacteria naturally die and nutrients naturally decay over the course of the modeling. These decays are linear with respect to the instantaneous abundance of the bacteria or the nutrient.

6. The resource-explicit growth of *E. coli* is based on the product of the two Monod terms, and not the minimum of the two as in Hammarlund et al. (2019).

Model Implementation & Parameterization: Twenty locations on the 1D line were selected randomly to seed *E. coli* (and *S. enterica*, if in the mutualistic coculture) colonies in the form of biomass at the initial time point, and nutrient was distributed evenly on the line as well. All variables in the equations above take values in Supplementary Table 1, which our research group together curated based on the previous ODE model describing the *E. coli* and *S. enterica* growth in our cross-feeding system (e.g. Hammarlund et al.; 2019; Hammarlund et al., 2021). The diffusion constants of nutrients were estimated based on another previous work in our group (Chacón et al., 2018). The initial colony biomass was set at 20 locations to be $E\left(t = 0, x_i\right) = 100$ and $S\left(t = 0, x_j\right) = 100$ for *E. coli* and *S. enterica*, respectively, at locations $x_i$ or $x_j$.

Note that very little (but non-zero) abundance of methionine and acetate were seeded evenly on the 1D line in the PDE model for the cross-feeding coculture. This follows previous convention in modeling the growth of an obligate cross-feeding mutualism (Harcombe et al., 2014; Harmmarlund et al., 2019; Harmmarlund et al., 2021). Briefly, the initial lactose was set to be $L\,(t\ = 0,x) = 1000\,\forall x$, methionine and acetate was seeded at $M\,(t\ = 0,x) = 0.01$ and $A\,(t\ = 0,x) = 0.01$ for all $x$ on the entire 1D line. Similarly in the monoculture model, the initial lactose and methionine concentrations were $L\,(t\ = 0,x) = M\,(t\ = 0,x) = 1000$ for all $x$. No acetate or *S. enterica* was present in the monoculture model.

Then, we non-dimensionalized the system in Eq. [S1] and [S2] as in Supplementary Discussion 2. We numerically solved the non-dimensionalized system Eq. [S3~S4] using MATLAB R2021b with the *pdepe* function. Finally, we calculated the growth rates and lag time of individual colonies by fitting a log-linear growth curve to the log-transformed biomass using in-house code in R v.3.3.3 (R Core Team, 2016).

**4. ODE-based antibiotic killing in populations with persisters**

*Model Setup & Implementation:* We let the unitless fractions of the non-persisters and persisters in a population of bacteria be denoted by constants $n$ and $p$, respectively. We first calculated the MDK99 measurements of the population in ampicillin killing as $p$ changes. To do so, we first measured the persister fraction, $p$, of the WT bacterial population experimentally (Materials & Methods), and then calculated the death rates of the non-persisters and persisters by taking the slopes of the respective log-transformed death phases with respect to the drug treatment time, $t$. The death rates were denoted by $\kappa_n < 0$ and $\kappa_p < 0$, respectively. Note that we always have $\kappa_n < \kappa_p$, given that persisters die more slowly than non-persisters. Afterwards, we simulated the kill curve by ampicillin in a population with persister, $p$, with a set of simple ODEs:

$$\frac{dN}{dt} = \kappa_n N,$$
$$\frac{dP}{dt} = \kappa_p P, \qquad \text{[S5]}$$

for the dynamics of survived non-persisters ($N$) and survived persisters ($P$) over time $t$.

To simulate antibiotic killing dynamics, we took $\kappa_n = -\,4.2829$ and $\kappa_p = -\,0.3233$ by fitting kill curves of the non-persisters and persisters in the experimental antibiotic killing assay for the monoculture *E. coli* on agar (Fig. 1C-D). With various initial time points $N_0 \equiv n$ and $P_0 \equiv p$, we analytically solved the system in Eq. [S5], and plotted $(N + P)$ with respect to $t$ for several total persister fractions $p$ (Fig. S2B). Then, we plotted the relationship between the measured MDK99 and $p$ in a population with susceptible (WT) cells and persisters at different $p$, shown by the function MDK99 $= f(p)$ in Fig. S2C.

#### Supplementary Discussion

##### 1. Determining cell-cell distance in shaken liquid and on agar surfaces

Here, we determined the average cell-cell distance on agar surfaces (in liquid) by assuming that cells are uniformly distributed in the 2D (3D) space. Each cell is assumed to be at the center of a very small square (lattice) with equal size. And the average cell-cell distance is estimated to be the size of the square (lattice).

We showed that the average cell-to-cell distances in shaken liquid ($5 \times 10^7$ CFU/mL) and on the nitrocellulose membrane surfaces ($5 \times 10^6$ CFU/membrane) are comparable (Fig. S4B):

- Monoculture on Membranes: $5 \times 10^6$ *E. coli* CFU/membrane ($OD_{600}$=0.005 for 2mL; average cell-to-cell distance: 37.2 $\mu$m) on each membrane to be placed on Hypho minimal agar.
- Monoculture in Liquid: $5 \times 10^7$ *E. coli* CFU/mL ($OD_{600}$=0.01 $mL^{-1}$; average cell-to-cell distance: 54.2 $\mu$m) in 5 mL shaken Hypho liquid minimal medium.
- Mutualism on Membranes: $5 \times 10^6$ CFU/membrane cells for both species ($OD_{600}$=0.005 for 2mL for *E. coli* and $OD_{600}$=0.0025 for 2mL for *S. enterica*; average cell-to-cell distance: 26.3 $\mu$m) on each membrane.
- Mutualism in Liquid: $5 \times 10^7$ CFU/mL for both species ($OD_{600}$=0.01 $mL^{-1}$ for *E. coli* and $OD_{600}$=0.005 $mL^{-1}$ *S. enterica*; average cell-to-cell distance: 43.1 $\mu$m) in 5 mL shaken Hypho liquid medium.

*Monoculture in Shaken Liquid:*

In shaken liquid, using $OD_{600}$=0.01/mL led to a cell density of $\rho_L = 5 \times 10^7$ CFU/mL for a cell number of $N_L = 5 \times 10^7$ in a volume of $V = 1$ $cm^3$. Assuming that cells were uniformly distributed, this means that each cell should be at the center of a very small square lattice with edge length, $x_{LM}$, and volume $V_L$. As a result, we estimate that individual volume of the small lattice is $V_L = \frac{V}{N_L} = \frac{1}{5 \times 10^7} = 2 \times 10^{-8}$ $cm^3$. Then the average distance between cells is $x_{LM} = 27.1$ $\mu$m.

*Monoculture on Surfaces:*

The diameter of each nitrocellulose membrane was $d = 4.7$ cm, so the area for each membrane is $A_M = 17.34$ $cm^2$. Using a density of ~$5 \times 10^6$ CFU/membrane for each species. This gives a cell density of $\rho_M = 2.88 \times 10^5$ cells/$cm^2$. Assuming that all cells were uniformly distributed, we estimate that each cell is at the center of a very small square with edge length, $x_{MM}$, and area $A_M$. As above, we estimate that $A_M = \frac{1}{2.88 \times 10^5} = 3.47 \times 10^{-6}$ $cm^2$. Then the average distance between cells is $x_{MM} = 18.6$ $\mu$m.

*Mutualism in Liquid:*

In shaken liquid, using $\rho_L = 5 \times 10^7$ CFU/mL for both species yielded a total cell number of $N_{LC} = 1 \times 10^8$ in a volume of $V = 1$ $cm^3$. Assuming that cells were uniformly distributed, this means that each cell should be at the center of a very small square lattice with edge length, $x_{LC}$, and volume $V_{LC}$. As a result, we estimate that individual volume of the small lattice is $V_{LC} = \frac{V}{N_{LC}} = \frac{1}{1 \times 10^8} = 1 \times 10^{-8}$ $cm^3$. Then the average distance between cells is $x_{LC} = 21.5$ $\mu$m.

*Mutualism on Surfaces:*

The diameter of each nitrocellulose membrane was $d = 4.7$ cm, so the area for each membrane is $A_M = 17.34$ $cm^2$. Using a density of ~$5 \times 10^6$ CFU/membrane for each species, we have a total cell density of $\rho_{MC} = 5.77 \times 10^5$ cells/$cm^2$ because we have two species. Assuming that all cells were uniformly distributed, we estimate that each cell is at the center of a very small square with edge length, $x_{MC}$, and area $A_{MC}$. As above, we estimate that $A_{MC} = \frac{1}{5.77 \times 10^5} = 1.734 \times 10^{-6}$ cm². The average distance between cells is then $x_{MC} = 13.2\ \mu$m.

Clearly, the average cell-to-cell distances in the four conditions above were all comparable.

**2. Nondimensionalization of the PDE model**

We performed nondimensionalization for both sets of equations to make all variables unit- and dimension-less. First, I non-dimensionalize the PDE systems in Eqs. [S1~S2] by defining $y = \frac{x}{Q}, \tau = t_0 t$, $u = \frac{E}{W}$, $s = \frac{S}{W}$, $m = \frac{M}{W}$, $a = \frac{A}{W}$, $l = \frac{L}{W}$, $\delta_m^2 = \frac{D_M}{t_0 Q^2}$, $\delta_a^2 = \frac{D_A}{t_0 Q^2}$, $\delta_l^2 = \frac{D_L}{t_0 Q^2}$, and $k_i = \frac{K_i}{W}$ for $i \in \{M, L, A\}$. Here, I let $t_0 = t^{-1}$ , $Q = 1$ unit of distance, and $W = 1$ cell unit per mL as in Hammarlund et al. (2021). Therefore, the non-dimensionalized, cross-feeding system in Eq. [S1], becomes:

$$\frac{\partial u}{\partial \tau} = \frac{r_E}{t_0}\left(\frac{m}{m + k_M}\right)\left(\frac{l}{l + k_L}\right)u - \frac{\kappa_E}{t_0}u, \qquad \text{[S3a]}$$

$$\frac{\partial s}{\partial \tau} = \frac{r_S}{t_0}\left(\frac{a}{a + k_A}\right)s - \frac{\kappa_S}{t_0}s, \qquad \text{[S3b]}$$

$$\frac{\partial m}{\partial \tau} = \delta_m^2 \nabla^2 m + \frac{p_M r_S}{t_0}\left(\frac{a}{a + k_A}\right)s - \frac{c_E}{t_0}\left(\frac{m}{m + k_M}\right)\left(\frac{l}{l + k_L}\right)u - \frac{\kappa_M}{t_0}m, \qquad \text{[S3c]}$$

$$\frac{\partial a}{\partial \tau} = \delta_a^2 \nabla^2 a + \frac{p_A r_E}{t_0}\left(\frac{m}{m + k_m}\right)\left(\frac{l}{l + k_l}\right)u - \frac{c_S}{t_0}\left(\frac{a}{a + k_a}\right)s - \frac{\kappa_A}{t_0}a, \qquad \text{[S3d]}$$

$$\frac{\partial l}{\partial \tau} = \delta_l^2 \nabla^2 l - \frac{c_L}{t_0}\left(\frac{m}{m + k_M}\right)\left(\frac{l}{l + k_L}\right)u - \frac{\kappa_L}{t_0}l. \qquad \text{[S3e]}$$

The boundary conditions become:

$$\frac{\partial n}{\partial y} = 0 \text{ at } y = 0,\ 5, \text{ for } n \in \{u, s, m, a, l\}. \qquad \text{[S3f]}$$

Similarly, the non-dimensionalized monoculture system of E. coli in Eq. [2] becomes:

$$\frac{\partial m}{\partial \tau} = \delta_m^2 \nabla^2 m - \frac{c_E}{t_0}\left(\frac{m}{m + k_M}\right)\left(\frac{l}{l + k_L}\right)u - \frac{\kappa_M}{t_0}m, \qquad \text{[S4a]}$$

$$\frac{\partial u}{\partial \tau} = \frac{r_E}{t_0}\left(\frac{m}{m + k_M}\right)\left(\frac{l}{l + k_L}\right)u - \frac{\kappa_E}{t_0}u, \qquad \text{[S4b]}$$

$$\frac{\partial l}{\partial \tau} = \delta_l^2 \nabla^2 l - \frac{c_L}{t_0}\left(\frac{m}{m + k_M}\right)\left(\frac{l}{l + k_L}\right)u - \frac{\kappa_L}{t_0}l, \qquad \text{[S4c]}$$

with similar boundary conditions:

$$\frac{\partial n}{\partial y} = 0 \text{ at } x = 0,\ 5, \text{ for } n \in \{u, m, l\}. \qquad \text{[S4d]}$$

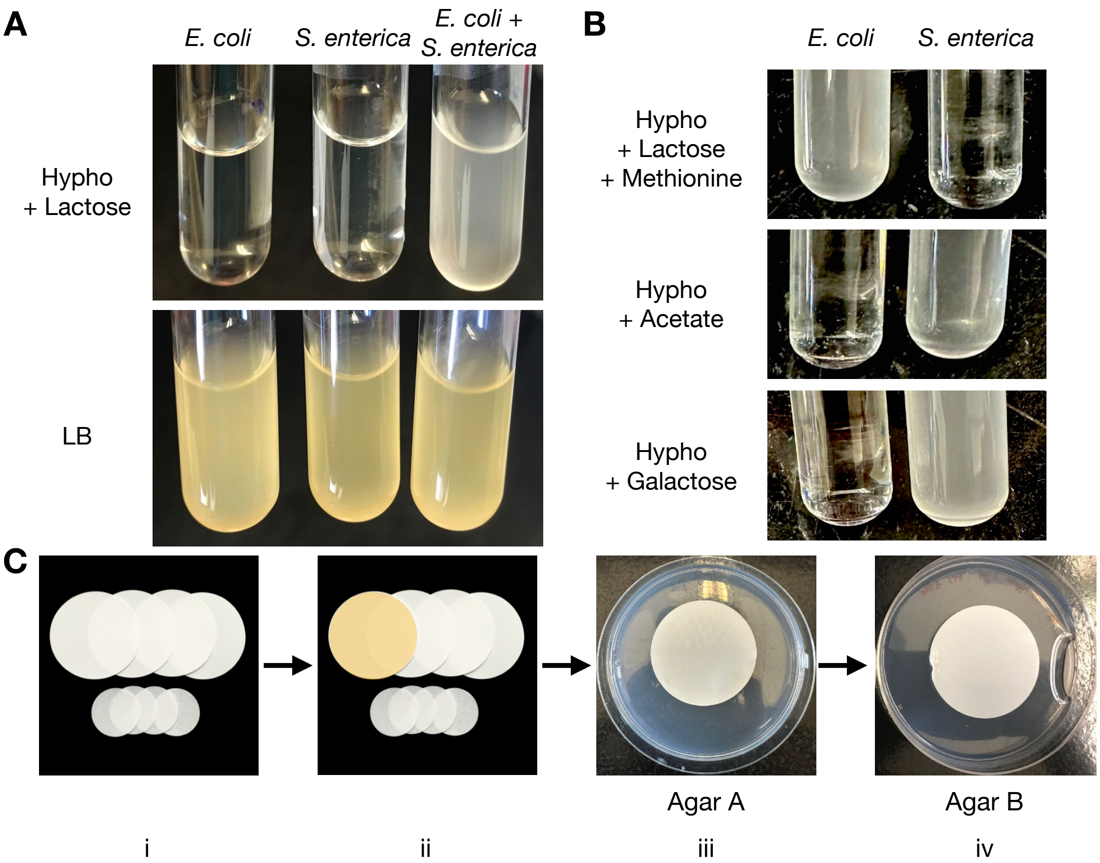


**Figure S1. An obligate microbial mutualism between *Escherichia coli ΔmetB* and the methionine-overproducer *S. enterica* in the lactose minimal media.** **A** Obligate mutualism is observed between the two domesticated *E. coli* and *S. enterica* strains (Harcombe, 2010; Harcombe et al., 2014) in the Hypho minimal medium with lactose as the only carbon source. Both species can be recovered in LB media. **B** The *E. coli* and *S. enterica* can also be studied in their respective monocultures while maintaining the same physiology as in the mutualistic coculture. In monoculture, *E. coli* can grow in Hypho minimal media supplemented with lactose and methionine, whereas *S. enterica* can grow in Hypho minimal media supplemented with acetate or galactose, which are the carbon wastes we believe *E. coli* is providing in the mutualistic coculture (Harcombe, 2010; Harcombe et al., 2018). Note that either species cannot grow alone in the other media due to auxotrophy. **C** Experimental setup in this work. On sterile nitrocellulose filter membranes (i), bacteria can be randomly distributed and immobilized by running a washed and diluted liquid culture through a membrane on a sterile funnel (ii). Then the membrane with bacteria (shade in orange) can be moved among multiple agar plates [e.g. including Agar A (iii) and Agar B (iv)]. Image of membranes in (i) and (ii) were retrieved from the manufacturer website.

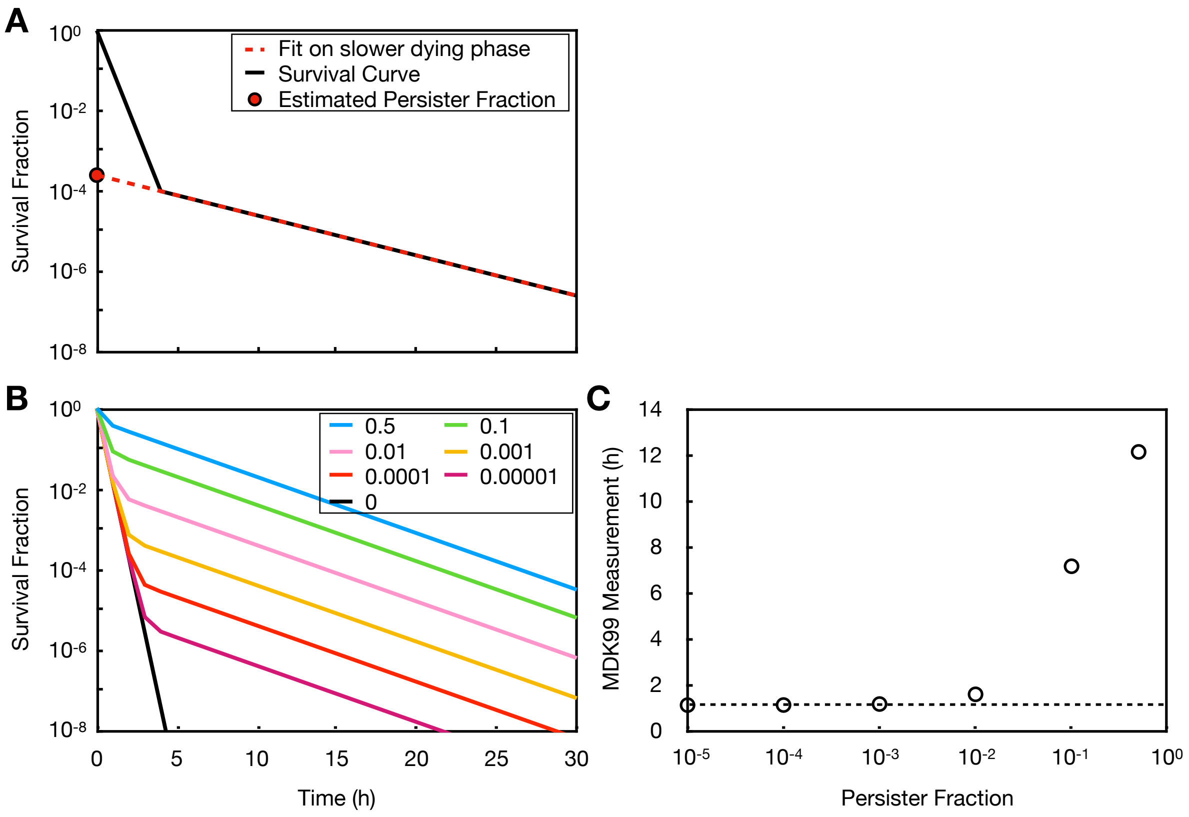


**Figure S2. Measuring antibiotic tolerance and persistence. A** For an experimentally measured survival curve over time (black line), we applied an exponential line (red dashed line) to fit the slower-dying phase of the survival curve, and then the intersection (red dot) between the fitted line and the *y*-axis is considered the initial persister fraction (Balaban et al., 2019). **B-C** Tolerance was measured as the "Standardized MDK99" with consideration of persister frequency. By definition, the regular MDK99 metric becomes inaccurate in measuring non-persisters' antibiotic tolerance when persister frequency becomes higher than 0.1%. **B** A simple ordinary differential equation (ODE) model (Supplementary Methods 4) was run to portray antibiotic killing of bacterial populations with persisters dying 10-times more slowly than the non-persisters. The model assumes no transition between persisters and non-persisters. Survival curves were simulated with different persister fractions at $t$=0 (color and legend). **C** The persister fraction affects MDK99's ability to portray antibiotic tolerance. Direct MDK99 calculations were made using the simulated data as in **A**. Note that no change was made to the non-persisters' tolerance in these simulations. Clearly, when persister fractions are above 0.1%, the MDK99 metric was inflated and fails to accurately measure the non-persisters' antibiotic tolerance.

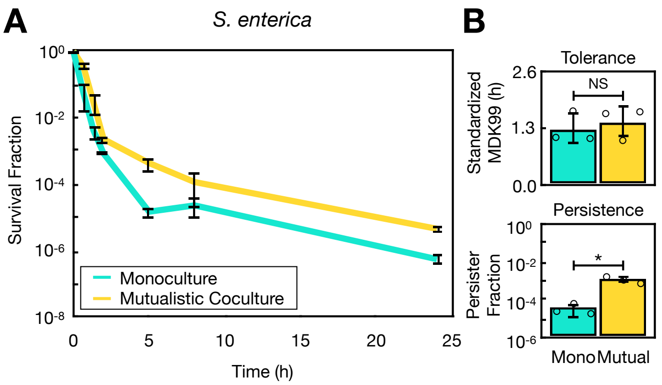


**Figure S3. Heightened antibiotic persistence phenotype was also observed in *S. enterica* in the spatially-structured mutualism. A** In the same experiment as in Fig. 1B-D, *S. enterica* shows differential survival curves in the antibiotic treatment between the mutualistic coculture and the monoculture media with galactose. **B** Using the same method as in Fig. 1B and S1B-C, we found that *S. enterica* has similar antibiotic tolerance between monoculture (Mono) and the mutualistic coculture (Mutual) (One-way ANOVA, $p$=0.6), but has ~32-fold higher persister frequency in mutualism than in monoculture (One-way ANOVA, $p$=0.01).

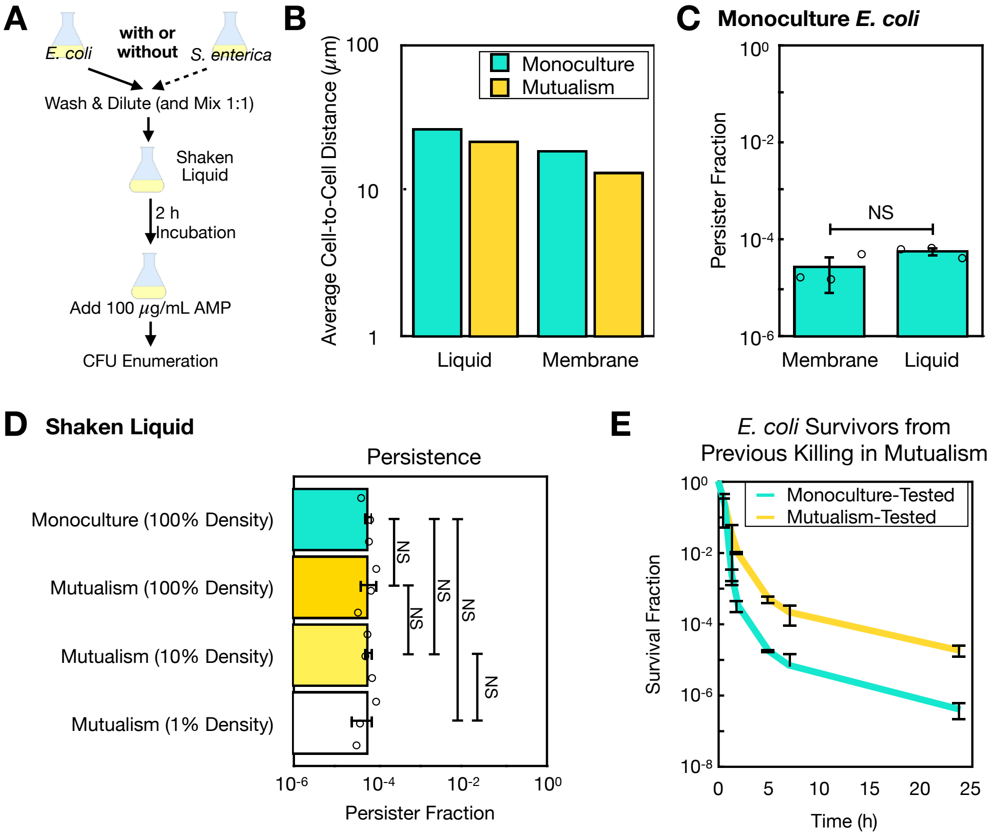


**Figure S4. Cross-feeding alone, spatial structure alone, the coculture density alone, or the spontaneous mutation alone was NOT sufficient to contribute to high persistence in *E. coli*. A** To test whether cross-feeding alone was sufficient to cause high *E. coli* persistence, we repeated the experiment in Fig. 1B in shaken liquid. **B** In all scenarios tested in this work, the average *E. coli* cell-cell distance was on the same order of magnitude. We assumed that on membranes (in liquid), each bacterial cell is at the center of a small square (cube) with identical edge length, with all of the squares (cubes) occupying the entire nitrocellulose membrane surface (liquid culture volume). Then the average cell-cell distance was then calculated as the distance between the center points of two adjacent squares (cubes) (Supplementary Discussion 1). **C** Spatial structure alone was NOT sufficient either to cause high antibiotic persistence in *E. coli* in monoculture. The log-phase monoculture *E. coli* shared a similar persister frequency in on nitrocellulose membranes and in shaken liquid (One-way ANOVA, $p$=0.1). **D** Coculture density alone was NOT sufficient to affect *E. coli* persistence**.** To test whether cell densities in the mutualistic cocultures affect *E. coli* persistence, we measured *E. coli* persister frequency at different starting densities in shaken liquid while maintaining the *S. enterica* : *E. coli* ratio to be 1:1. We found that lowering the *E. coli* density down to 1% of our original experiment (Fig. 1B) was insufficient to change persister frequency (One-way ANOVA, Tukey's HSD $p$>0.9 for all pairwise comparisons). **E** The heightened persistence in the spatially-structured mutualistic coculture was NOT due to spontaneous mutations. In our structured membrane setup, we almost always used the *E. coli* cell density at 5E6 CFU/membrane that limits potential for spontaneous mutations to increase *E. coli*'s ampicillin resistance. To more rigorously confirm this, we isolated one survived colony from each of the three biologically independent mutualistic coculture killing experiment on membranes (Fig. 1C), and repeated the killing experiment (Fig. 1B-C) to them. The resulting survival curves were similar to that as in Fig. 1C.

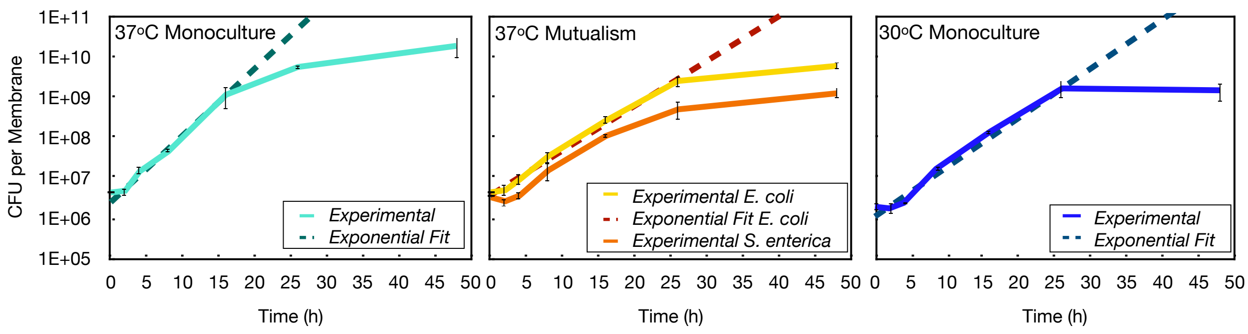


**Figure S5. *E. coli* growth curves on membranes.** The *E. coli* growth curves were plotted on nitrocellulose membranes in various growth conditions. Cells were immobilized on nitrocellulose membranes as in Fig. 1B, and incubated statically at 37 ºC or 30 ºC as specified in the figure. At 2, 4, 8, 16, 26, 48 hours, membranes from each biological replicate were taken off from the agar and rinsed in 5 mL saline for CFU enumeration (disruptive sampling). The dotted lines were lines of best fit for the log-phase of the growth curves. The growth rates were the slopes of these lines, and the lag time was the *x*-axis values at the intersection between the log fit and the initial CFU value (Materials & Methods).

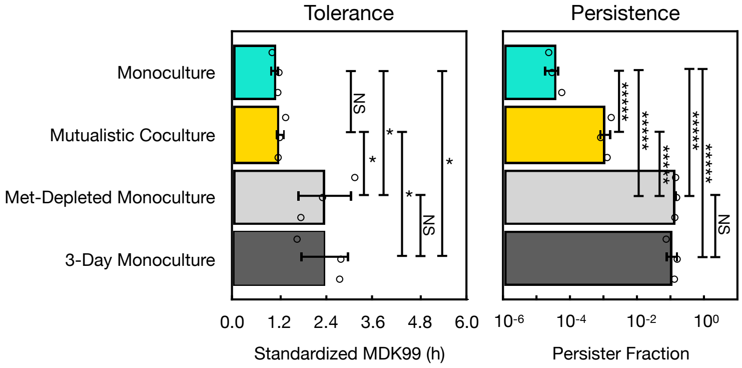


**Figure S6. Starvation in *E. coli* on agar increased antibiotic tolerance and persistence.** We tested whether the starvation for methionine was sufficient to increase antibiotic persistence. Thus, we treated *E. coli* with ampicillin as in Fig. 1B on the methionine-depleted Hypho agar with lactose, and found that such starvation increased persistence (One-way ANOVA, Tukey's HSD $p$<1E-7) as well as tolerance (One-way ANOVA, Tukey's HSD $p$=0.04) compared to the monoculture with methionine supplementation. This is different than the effect of growing in the cross-feeding mutualism, which only changed persistence (One-way ANOVA, Tukey's HSD $p$=8E-6) but not tolerance (One-way ANOVA, Tukey's HSD $p$>0.9). Similarly, we starved *E. coli* for 3 days in the monoculture media, and found that it gains significantly higher tolerance (One-way ANOVA, Tukey's HSD $p$= 0.04) and persistence (One-way ANOVA, Tukey's HSD $p$<1E-7) than its log-phase counterpart. Note that the experiments for Fig. 1F, 2B, and S6 were performed together.

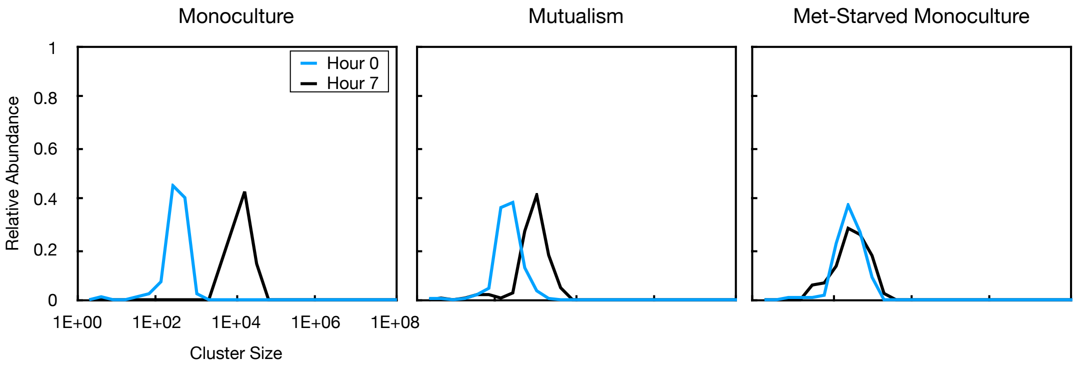


**Figure S7. Tracking biomass for single *E. coli* cell clusters.** Consistent with Levin-Reisman et al. (2010) and Fridman et al. (2014), we measured lag time as the time it takes for a single CFP-labeled *E. coli* cluster identified at Hour 0 to gain biomass till a certain threshold (in our case, to gain 10% of biomass). Consistent with our intuition, we found that by Hour 7, the *E. coli* clusters on average gain biomass by 10~100 fold in the mutualistic coculture with *S. enterica* or the monoculture, but almost no biomass was gained in the methionine- (Met-) starved monoculture.

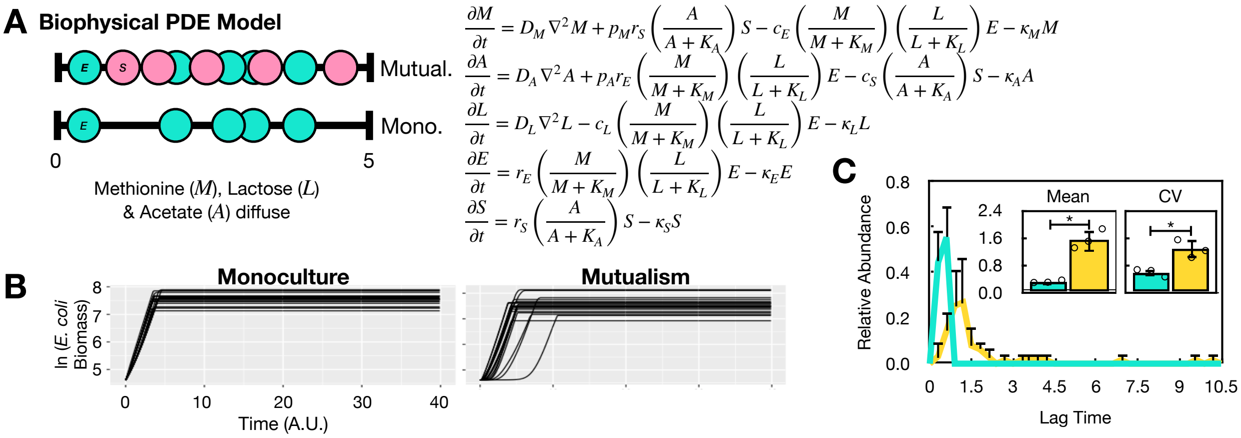


**Figure S8. A simple mathematical model supports the microscopy experiment results**. **A** A biophysical (resource-consumption, reaction-diffusion) PDE model was designed to describe *E. coli* (cyan) growth at a single colony level on a 1-dimension world with and without *S. enterica* (pink). Briefly, lactose is initially set up in the entire 1-dimension line. The *E. coli* (*E*) and *S. enterica* (*S*) cells were seeded at 20 random locations, and the cells can grow (increase biomass) by consuming nutrient. *E. coli* consumes lactose (*L*) and methionine (*M*) to grow, while *S. enterica* consumes acetate (*A*). *E. coli* secretes acetate as it grows, and *S. enterica* secretes methionine. The nutrient diffuses in the world at diffusion constants $D_i$ for $i = M, A, L$ that we chose based on Chacón et al. (2018). There is also a natural decay rate for all terms, $\kappa_i$ for $i = M, A, L, E, S$. See Supplementary Methods 3 for details. **B** When seeded at the same location in monoculture or in mutualism with *S. enterica*, the individual *E. coli* growth curves were very different. Results shown here are from one representative independent simulation from **A**. **C** Characterizing all growth curves in each simulation following Materials & Methods, we found significantly larger mean (One-way ANOVA, *p*=2E-3) in the *E. coli* lag time in the cross-feeding coculture than in monoculture, and this was mainly caused by a wider and right-shifted lag time distribution (coefficient of variation, or CV, was higher, one-way ANOVA, *p*=0.01).

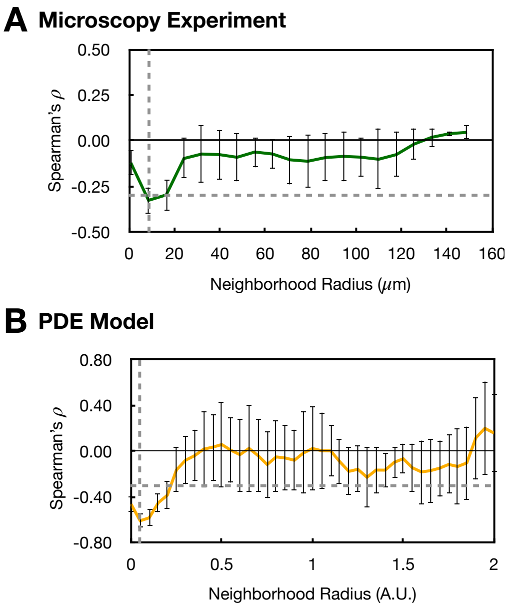


**Figure S9. *E. coli* and *S. enterica* typically interact in small neighborhoods.** Here we used a modified method from Dal Co et al. (2020) to measure the interaction range between *E. coli* and *S. enterica* in the mutualistic coculture. In brief, we calculated the initial *S. enterica* biomass fraction in neighborhoods with various sizes (radius), and correlated this biomass fraction at all radius sizes with the *E. coli* cell cluster lag time (Fig. 3B-C). Then, the neighborhood radius was plotted against the Spearman's $\rho$ in **A-B**. The horizontal gray dashed line denotes Spearman's $\rho$=-0.3. The vertical gray dashed line indicates the radius size where the correlation was best, which was defined as the interaction range (Dal Co et al., 2020). Error bars denote standard deviation of the average measurements from 3 biologically independent trials. **A-B** Both the microscopy experiment and the PDE model revealed that *E. coli* and *S. enterica* usually interact in very small neighborhoods (~10 $\mu$m in experiment). However, the correlation in the PDE model (Spearman's $\rho$=-0.610±0.0633) was much better than in the microscopy experiment ($\rho$=-0.327±0.0735, $p$<0.002).

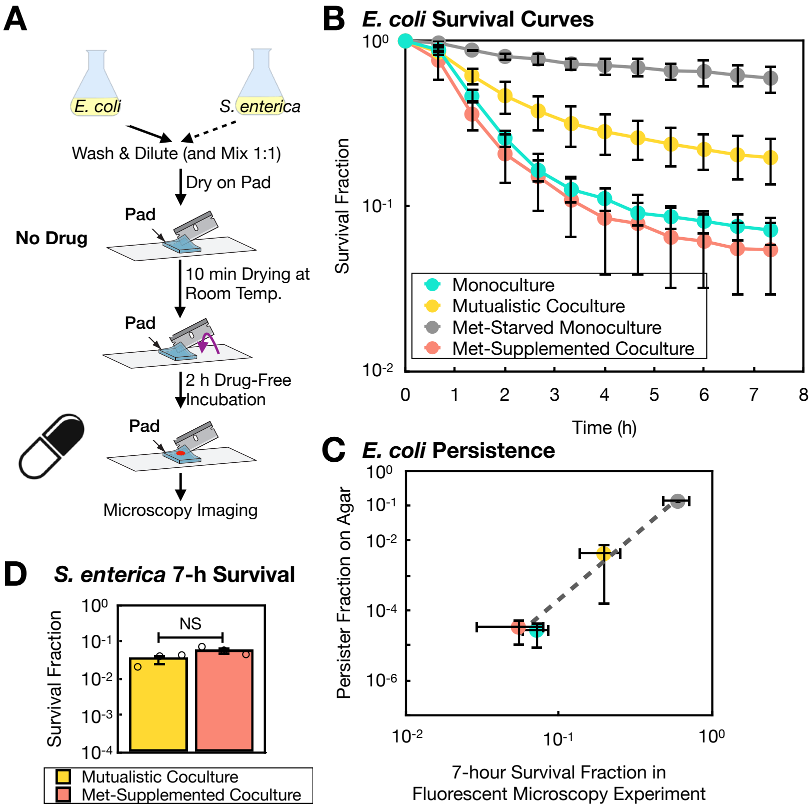


**Figure S10. Predicting the *E. coli* death time on a single-cell level. A** A fluorescent microscopy setup to track the *E. coli* ampicillin-induced death with spatial structure. In this panel, all illustrations that contain agarose pads were edited from the original one in Skinner et al. (2013). A 1.5 µL cell droplet of $OD_{600}$=0.05 from saline-washed *E. coli* and $OD_{600}$=0.025 of *S. enterica* was first incubated on top of the Hypho minimal agarose pad to dry at room temperature, and then flipped over (purple arrow) on a microscopic slide for incubation at 37 $^{o}$C. Note that we have tested that this concentration leads to similar cell density in the microscopic frame as on the nitrocellulose membrane setup. Afterwards, a 1.5 µL ampicillin droplet was added on top of the pad (red dot) such that the entire agarose pad will eventually reach 100 µg/mL. **B** Repeating the experiment in all four conditions in Fig. 1F and S6 on the microscopy setup recreated the survival curves for *E. coli* that shared the same trends. **C** The mean survival fraction data after a 7-hour ampicillin treatment under the agarose pad can predict the mean persister fraction measurements on nitrocellulose membranes in Fig. 1F and Fig. S6 (Adjusted $R^2$=0.9673, $p$=0.01). **D** The mutualistic (yellow) and the methionine-supplemented (red) cocultures led to similar *S. enterica* responses to the ampicillin treatment. We found that the differences in the 7-hour survival fractions in *S. enterica* between the two conditions were borderline different ($p$=0.09).

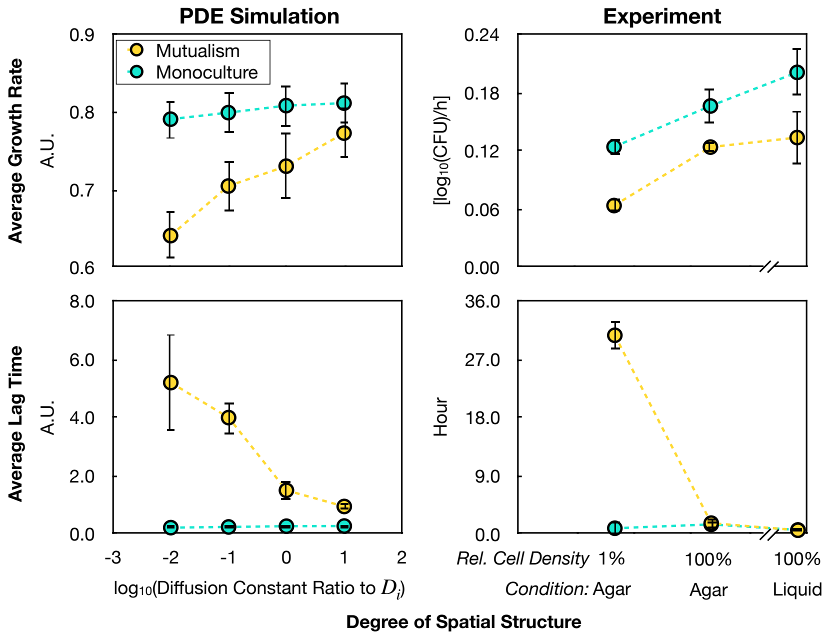


**Figure S11. Choosing microscopy over the ScanLag approach to study sub-population growth of *E. coli*.** ScanLag is a method where the growth of bacteria is tracked on a single-colony level on agar plates on an office scanner that periodically takes images of the plate (Levin-Reisman et al., 2010). This established method has been successfully implemented in numerous studies (e.g. Fridman et al., 2014; Levin-Reisman et al., 2017; Chacón et al., 2018; Vulin et al., 2018) to understand individual heterogeneity of a given bacterial population. We decided to perform fluorescent microscopy rather than this method because this method was shown (Levin-Reisman et al., 2010; Fridman et al., 2014) to work best with 100~200 colonies on an agar plate with a diameter of 10 cm. This density is about $10^{-5}$ times less than on the nitrocellulose membrane setting in the current work. Both the PDE model and our experiments show that lowering cell density will artificially augment the mean of the lag time for *E. coli*.

| Parameter | Unit | Value | Biological Interpretation | Source |
|---|---|---|---|---|
| $x$ | Arbitrary unit of distance | 1 | Unit of space. | Definition |
| $t$ | Arbitrary unit of time | 1 | Unit of time. | Definition |
| $Y$ | Arbitrary unit of distance | 5 | Total size of the 1-dimension modeling canvas. | Definition |
| $E$, $S$, $M$, $A$, $L$ | Cell unit/mL | - | *E. coli*, *S. enterica*, methionine, acetate, and lactose, respectively. | Definition; Hammarlund et al. (2021) |
| $D_M$ | $x^2/t$ | 0.01 | Diffusion constant of methionine. | Estimated based on Chacón et al. (2018) |
| $D_A$, $D_L$ | $x^2/t$ | 0.05 | Diffusion constants of the acetate and lactose sugars. | Estimated based on Chacón et al. (2018) |
| $p_M$ | Unitless | 1.56 | Production rate of methionine by *S. enterica*. | Hammarlund et al. (2021); adjusted with unpublished laboratory data |
| $p_A$ | Unitless | 1.01 | Production rate of acetate by *E. coli*. | Hammarlund et al. (2021); adjusted with unpublished laboratory data |
| $c_E$ | $t^{-1}$ | 0.1 | *E. coli* consumption rate of methionine. | Hammarlund et al. (2021); adjusted with unpublished laboratory data |
| $c_L$ | $t^{-1}$ | 1.0 | *E. coli* consumption rate of lactose. | Hammarlund et al. (2021); adjusted with unpublished laboratory data |
| $c_S$ | $t^{-1}$ | 1 | *S. enterica* consumption rate of acetate. | Hammarlund et al. (2021); adjusted with unpublished laboratory data |
| $r_E$ | $t^{-1}$ | 1 | *E. coli* maximum growth rate. | Hammarlund et al. (2021) |
| $r_S$ | $t^{-1}$ | 0.5 | *S. enterica* maximum growth rate. | Hammarlund et al. (2021) |
| $K_M$, $K_A$, $K_L$ | Cell unit/mL | 1 | Half-saturation methionine, acetate, lactose concentration for bacterial growth. | Hammarlund et al. (2021); adjusted for simplicity |
| $\kappa_E$, $\kappa_S$, $\kappa_M$, $\kappa_A$, $\kappa_L$ | $t^{-1}$ | 5E-09 | Natural decay rate of *E. coli*, *S. enterica*, methionine, acetate, and lactose. | Estimated and adjusted for simplicity |

**Supplementary Table 1. Parameters Used in the PDE Model.**
